# Sphingosine-1-phosphate receptors 1 and 3 regulate the expression of scavenger receptor B1 in human aortic endothelial cells

**DOI:** 10.1101/2020.04.23.058263

**Authors:** Dongdong Wang, Lucia Rohrer, Arnold von Eckardstein

## Abstract

Several vasoprotective functions of high-density lipoproteins (HDL) on the endothelium have been shown to depend on the presence of sphingosine-1-phosphate (S1P) receptors (S1PRs) as well as scavenger receptor class B type 1 (SR-B1). Interference with the presence of S1P or the activity of S1PR1 or S1PR3 mimics many effects seen by the interference with SR-B1. This raises the question on interactions between S1P receptors and SR-B1. We investigated the influence of S1PRs on SR-B1 expression in human aortic endothelial cells. Silencing or pharmacological inhibition of S1PR1 or S1PR3 down-regulated *SCARB1* mRNA expression as well as SR-B1 protein abundance. RNA interference with S1PR1 or S1PR3 also decreased cellular association of ^125^I-HDL with HAECs. Further mechanistic studies showed that knockdown of S1PR1 or S1PR3 reduced SR-B1 protein by inducing its degradation through deceasing Akt activity. Moreover, silencing of S1PR1 or S1PR3 suppressed *SCARB1* mRNA expression by decreasing cellular cAMP levels. In conclusion, we provide evidence for an as yet unappreciated interaction, namely the regulation of SR-B1 abundance by S1PRs on both transcriptional and post-translational levels, suggesting that interactions of S1PRs and SR-B1 regulate signaling functions of HDL as well as uptake of lipoproteins in endothelial cells.

## INTRODUCTION

Scavenger receptor class B type 1 (SR-B1) is a member of cluster of differentiation 36 (CD36) superfamily (1). The SR-B1 encoding *SCARB1* gene is expressed in many tissues and cell types, including liver, steroidogenic tissues, macrophages, and endothelial cells (2-4). SR-B1 is a multifunctional receptor which is involved in the pathogenesis of atherosclerosis but also other diseases including cancer (2, 3, 5). The prototype function is the mediation of cholesterol flux into and out of cells (1, 3). SR-B1 mediates the selective uptake of cholesteryl esters mostly from high-density lipoproteins (HDL) into liver, adrenals, testes, or ovaries which use cholesterol for the formation of bile and steroid hormones, respectively (1, 3, 6, 7). From macrophages, endothelial and other cells, SR-B1 rather mediates cholesterol efflux to HDL. The relative activity of cholesteryl ester consuming and producing enzymes in the cell and the plasma appear to dictate the directionality of the flux (1). The presence or absence of SR-B1, however, also affects other cellular functions either indirectly through changes in the distribution of cholesterol within the cell and cell membrane or directly by interaction with adapter proteins such as PDZK1 or DOCK4 (8-10). In endothelial cells, SR-B1 regulates the uptake of both LDL and HDL for transendothelial transport (9-13). SR-B1 in endothelial cells was also found to be a limiting factor for the ability of HDL to promote nitric oxide production as well as proliferation and migration, and to inhibit cellular adhesion molecule, expression and, as the consequence, leukocyte transmigration (10, 14-19). These vasoprotective and anti-inflammatory functions have also been assigned to sphingosine-1-phosphate (S1P) which is enriched in HDL through its binding to apolipoprotein M. S1P interacts with five different G-protein coupled S1P receptors (18, 20-24). They include S1PR1, S1PR2, and S1PR3 which are expressed in endothelial cells and regulate the development and function of the vasculature (20, 21). Interference with the presence of S1P or the activity of S1PR1 or S1PR3 mimics many effects seen by the interference with SR-B1. This raises the question on interactions between S1P receptors and SR-B1. For example, it has been suggested that SR-B1 tethers HDL on the cell surface and thereby increases the likelihood of S1P/S1P-receptor interaction (2, 25). We here investigated the influence of S1PRs on SR-B1 expression.

## MATERIALS AND METHODS

### Cell culture

Human aortic endothelial cells (HAECs) from Cell Applications Inc. (304-05a) were cultured in endothelial basal medium (EBM)-2 (LONZA Clonetics^™^ CC-3156) with 5% fetal bovine serum (GIBCO), supplemented with endothelial growth medium (EGM)-2 SingleQuots (LONZA Clonetics CC-4176) containing human epidermal growth factor (hEGF, CC-4317A), vascular endothelial growth factor (VEGF, CC-4114A), R3-insulin-like growth factor-1 (R3-IGF-1, CC-4115A), ascorbic acid (CC-4317A), hydrocortisone (CC-4112A), human fibroblast growth factor-beta (hFGF-β, CC-4113A), heparin (CC-4396A), and gentamicin/amphotericin-B (GA-1000, CC-4381A) at 37 °C in a humidified 5% CO2, 95% air incubator.

### Small interfering RNA (siRNA) transfection

Endothelial cells were reverse transfected with siRNA targeted to S1PR1 (Ambion: Cat. no. s4447, s4449; Dharmacon: Cat. no. M-003655-02-0005), S1PR3 (Ambion: Cat. no. s4453, s4455; Dharmacon: Cat. no. M-005208-02-0005), SR-B1 (Ambion: Cat. no. s2648, s2649), p38 mitogen-activated protein kinases (p38 MAPK, Ambion: Cat. no. s3585), extracellular signal-regulated kinase kinase (MEK, Ambion: Cat. no. s11167), Akt1 (Ambion: Cat. no. s659, s660), Akt2 (Ambion: Cat. no. s1215, s1217), Akt3 (Ambion: Cat. no. s19427, s19429), non-silencing control siRNA (Ambion: Cat. no. 4390843; Dharmacon: Cat. no. D-001810-10-50) at a final concentration of 10 or 20 nmol/L for 72 h using Lipofectamine® RNAiMAX reagent (Invitrogen, 13778-150) in an antibiotic-free EGM-2 medium (without GA-1000). The transfection procedure was performed according to the manufacturer’s transfection protocol. Efficiency of transfection was confirmed by using qRT-PCR and western blot analyses.

### Western blot analyses

Western blot analyses were performed as described previously (26). HAECs were seeded at a density of 0.4 × 10^6^ cells/well in 6-well plates. The cells were reverse transfected with siRNA targeted to SR-B1, S1PR1, S1PR3, p38 MAPK, MEK, Akt1, Akt2, or Akt3 for 72 h, or treated with FTY720 phosphate (FTY720-P, Cayman: 10008639) or triciribine (Sigma, T3830) as indicated. To examine a putative liver X receptor (LXR), retinoid X receptor (RXR) and cyclic AMP (cAMP) dependency, the LXR agonist TO901317 (10 μM), the RXR agonist bexarotene (1 μM) and cAMP were used. The cells were lysed in ice-cold RIPA buffer with cOmplete^™^ protease inhibitor (Roche) and/or phosphatase inhibitors (Sigma, P0044) for 10 min before centrifugation (20 817 g for 15 min) to remove cellular debris. Concentration of total cellular protein was measured according to the DC^™^ protein assay according to the manufacturer’s instruction (BIO-RAD).

Samples (20 μg total protein/sample) were loaded and separated *via* SDS-PAGE (10%), and transferred to a polyvinylidene fluoride (PVDF, GE Healthcare) membrane. After blocking for 1 h with 5% nonfat dry milk or BSA in PBS-Tween (PBST), membranes were incubated with the following primary antibodies at 4°C overnight: S1PR1 (Abcam: Cat. no. ab125074, 1:1000), S1PR2 (Abcam: Cat. no. ab125074, 1:500), S1PR3 (Abcam: Cat. no. ab108370, abcam, 1:1000), SR-B1 (Novus: Cat. no. NB400-131, 1:500), ABCG1 (Novus: Cat. no. NB400-132, 1:500), total Akt (Cell signalling: Cat. no. 9272s, 1:1000), phospho-Akt (Ser473) (Cell signalling: Cat. no. 9271, 1:1000), Akt1 (Cell signalling: Cat. no. 2938, 1:1000), Akt2 (Cell signalling: Cat. no. 3063, 1:1000), Akt3 (Cell signalling: Cat. no. 3788, 1:1000), MEK1/2 (Cell signalling: Cat. no. 8727, 1:2000), p38 MAPK (Cell signalling: Cat. no. 9212, 1:2000), or TATA-binding protein (TBP, Abcam: Cat. no. ab51841-100, 1:5000). After washing with PBST, membranes were incubated with polyclonal rabbit anti-goat immunoglobulins/HRP (Dako: Code no. P0449, 1:2000), polyclonal rabbit anti-mouse immunoglobulins/HRP (Dako: Code no. P0260, 1:1000), or polyclonal rabbit anti-rabbit immunoglobulins/HRP (Dako: Code no. P0448, 1:2000) at room temperature for 1 h. Membranes were further incubated with chemiluminescence substrate (SuperSignal West Femto Maximum Sensitivity Substrate, or SuperSignal™ West Pico PLUS Chemiluminescent Substrate) for 1 minute (Thermo scientific). Protein bands were visualized with the Fusion FX UILBER LOURMAT (Vilber), and quantified with ImageJ software.

To measure SR-B1 protein stability, HAECs were seeded in 6-well plates as described above, and then co-treated with triciribine (30 μM) and the protein synthesis inhibitor cycloheximide (CHX, 100 μM) for different time points (0, 1, 2, 3, 4, and 6 h). The protein expression was detected by Western blot analyses. GraphPad Prism was applied to calculate the protein half-life (t_1/2_).

### Quantitative reverse transcription-polymerase chain reaction (qRT-PCR)

HAECs were seeded at a density of 0.4 × 10^6^ cells/well in 6-well plates. Where indicated, cells were reverse transfected with siRNA targeted to SR-B1, S1PR1, S1PR3, p38 MAPK, MEK, Akt1, Akt2, or Akt3 for 72 h, or treated with FTY720-P or triciribine as indicated.

After transfection or drug treatment, total RNA was extracted from cells using TRI reagent (Sigma, T9424) according to the manufacturer’s instruction. Concentration of total RNA was measured with NanoDrop 1000 (Witec AG). Genomic DNA was removed by digestion using DNase I Recombinant (Roche, 04716728001) and RiboLock RNase Inhibitor (Thermo Scientific, MAN0012010) according to the manufacturers’ instructions. cDNA was synthesized from 1 μg of total RNA by using the RevertAid Reverse Transcriptase kit. LightCycler® 480 SYBR Green I Master kit was used for quantification of SR-B1, S1PR1, S1PR2, and S1PR3 mRNA expression *via* the LightCycler® 480 System (Roche). Relative mRNA levels were quantified with the ΔCT method, using human GDPDH as an endogenous control. Gene specific primers were listed as followed: *S1PR1* (Forward primer (For): GTC TGG AGT AGC GCC ACC; Reverse primer (Rev): GTA GTC AGA GAC CGA GCT GC), *S1PR3* (For: TGA TCG GGA TGT GCT GGC; Rev: GAG TAG AGG GGC AGG ATG GTA), *SCARB1* (For: CTG TGG GTG AGA TCA TGT GG; Rev: GCC AGA AGT CAA CCT TGC TC), *GAPDH* (For: CCC ATG TTC GTC ATG GGT GT; Rev: TGG TCA TGA GTC CTT CCA CGA TA).

### Cellular cAMP level assay

HAECs were seeded at a density of 1 × 10^4^ cells/well in 96-well plates. The cells were reverse transfected with siRNA targeted to S1PR1, S1PR3 for 72 h, and then treated with SEW2871 (Cayman Chemical, 50 nM), CYM5541 (Tocris Bioscience, 100 nM) or vehicle (0.1% DMSO) for 1h as indicated. The untreated cells were used as a control. cAMP-Glo™ Assay kit (Promega, Cat. no. V1501) was used to measure cellular cAMP levels *via* GloMax® Discover Microplate Reader (Promega, Cat. no. GM3000) according to the manufacturer’s instruction. The luminescence output of the assay was calculated by the change in relative luminescence (ΔRLU = RLU (untreated sample) – RLU (treated sample)). Using these ΔRLU values and the linear equation generated from the standard curve (cAMP), calculate the cellular cAMP concentrations.

### Cellular association of HDL

The radiolabeled HDL cellular association assay was performed in line with previously published studies (27). HDL (1.063 < density < 1.210 g/ml) was isolated from human plasma of blood healthy donors by sequential ultracentrifugation (28, 29). HDL was labeled with Na^125^I by using the McFarlane monochloride methods modified for lipoproteins (29, 30). Specific activities of radiolabeled HDL were between 200-800 cpm/ng of protein. HAECs were seeded at a density of 0.1 × 10^6^ cells/well in 24-well plates. Cells were reverse transfected with siRNA targeted to SR-B1, S1PR1 or S1PR3, or non-silencing control siRNA (NC siRNA) for 72 h as described above. After siRNA transfection, the cells were incubated with 10 μg/mL of ^125^I-HDL without (total) or with (unspecific) 400 μg/mL (40×) of non-labeled HDL in DMEM medium (25 mmol/L HEPES, 100 U/ml benzylpenicillin, 100 μg/ml streptomycin, and 0.2% BSA) for 1 h at 37°C for HDL cellular association experiments. Specific HDL cellular association = (The total value (cpm) - unspecific value (cpm))/protein content

### Statistical analysis

For determination of differences between two groups, a two-tailed unpaired Student’s *t*-test was applied after data were tested for normality. For multiple comparisons, data were analyzed by one-way ANOVA followed by Bonferroni test to compare means between groups. *P* < 0.05 was considered statistically significant. GraphPad Prism (Version 8.0.0, GraphPad Inc., La Jolla, CA) was used for statistical analysis and figure generation.

## RESULTS

### Down-regulation of S1PR1 or S1PR3 protein decreases SR-B1 expression in HAECs

On both the mRNA and protein level, we found HAECs to express S1PR1 and S1PR3 at significant amounts, whereas S1PR2 was hardly detectable (Supplemental Figures S1). RNA interference decreased the expression of both S1PR1 and S1PR3 by 79% to 85% on the mRNA level and by 49% to 51% on the protein level (Supplemental Figures S1). Knockdown of S1PR1 or S1PR3 decreased both SR-B1 protein and mRNA expression by 71% to 84% and 56% to 61%, respectively, compared to non-coding siRNA (Figures 1A and 1B). The respective percentages were higher than 90% for silencing of *SCARB1*. Furthermore, knockdown of S1PR1 or S1PR3 significantly decreased cellular association of ^125^I-HDL with HAECs by 18% to 19% compared to 28% upon knock-down of *SCARB1* (Figure 1C).

**Figure 1.**
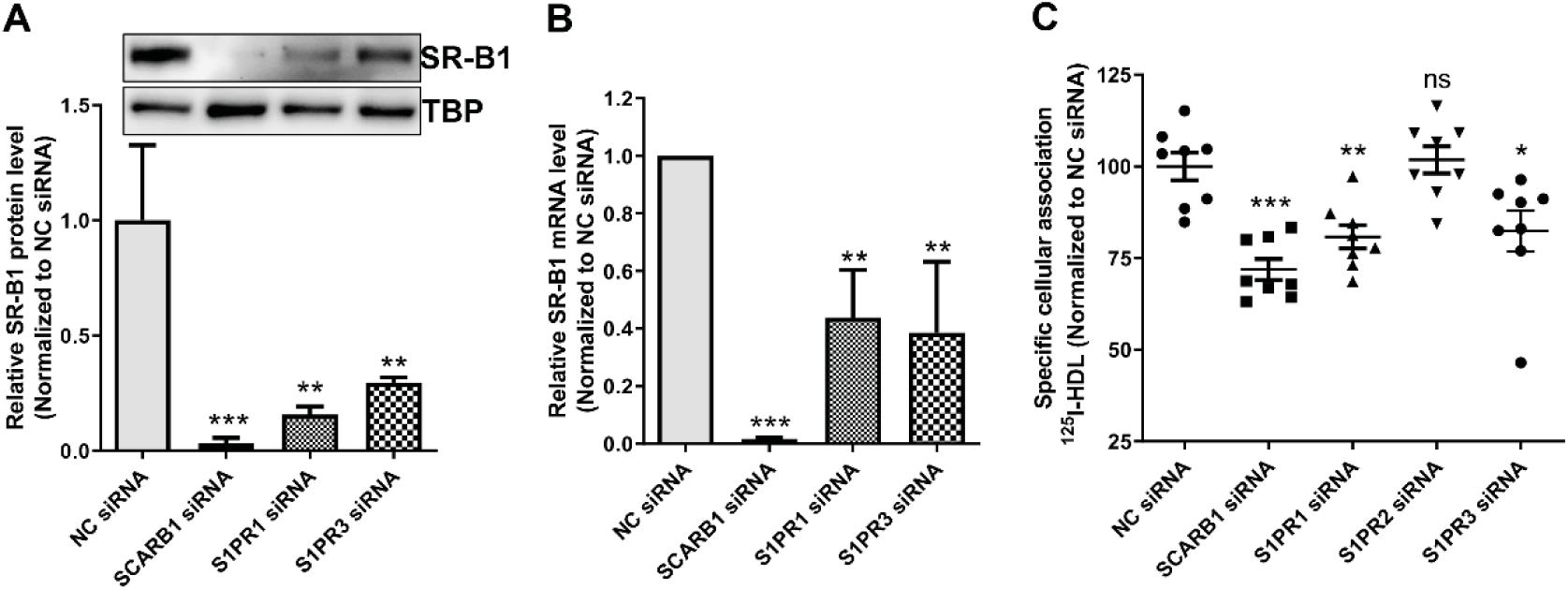
Knockdown of S1PR1 or S1PR3 protein decreases SR-B1 activity in HAECs. **A and B**. Knockdown of S1PR1 or S1PR3 dramatically decreases both SR-B1 protein and mRNA expression. HAECs were seeded at a density of 0.4 × 10^6^ cells/well in 6-well plates. The cells were reverse transfected with siRNA (10 nM) targeted to SR-B1, S1PR1 or S1PR3, or with non-silencing control siRNA (NC siRNA) for 72 h. The expression of SR-B1 protein and mRNA was determined by western blot analyses and qRT-PCR, respectively. TATA-binding protein (TBP) and GAPDH mRNA were used as the internal control for western blot analyses and RT-qPCR, respectively. **C**. Knockdown of S1PR1 or S1PR3 but not S1PR2 reduces endothelial HDL uptake. HAECs were seeded at a density of 0.1 × 10^6^ cells/well in 24-well plates. Cells were reverse transfected with siRNA (10 nM) targeted to SR-B1, S1PR1 or S1PR3, or with NC siRNA for 72 h. Cells were then incubated with 10 μg/mL of ^125^I-HDL for 1 h in the absence (total) or presence (unspecific) of 40-fold excess of unlabeled HDL. Specific HDL cellular association was calculated by subtracting unspecific values from total values. Data are shown as mean ± SD from three independent experiments, each performed in triplicate in case of the HDL cellular association assay (**C**). *P < 0.05, **P < 0.01, **P < 0.001, and ns not significant (Student’s *t*-test or ANOVA with Bonferroni test).

To confirm the effects of loss-of-S1PR-function on SR-B1 expression, we applied FTY720-P, which induces degradation of S1PRs during long term treatment of cells (31). Incubation with 10 nM FTY720-P down-regulated protein levels of both SR-B1 and S1PR1 in a time dependent manner by up to 38% and 27%, respectively (Figure 2A and 2B). During 24 h incubation, the suppression of SR-B1 mRNA by FTY720-P was dose dependent with a maximum of 56% seen at 100 nM (Figure 2C). The 43% suppression of SR-B1 protein however was already maximal at 10 nM (Figure 2D).

**Figure 2.**
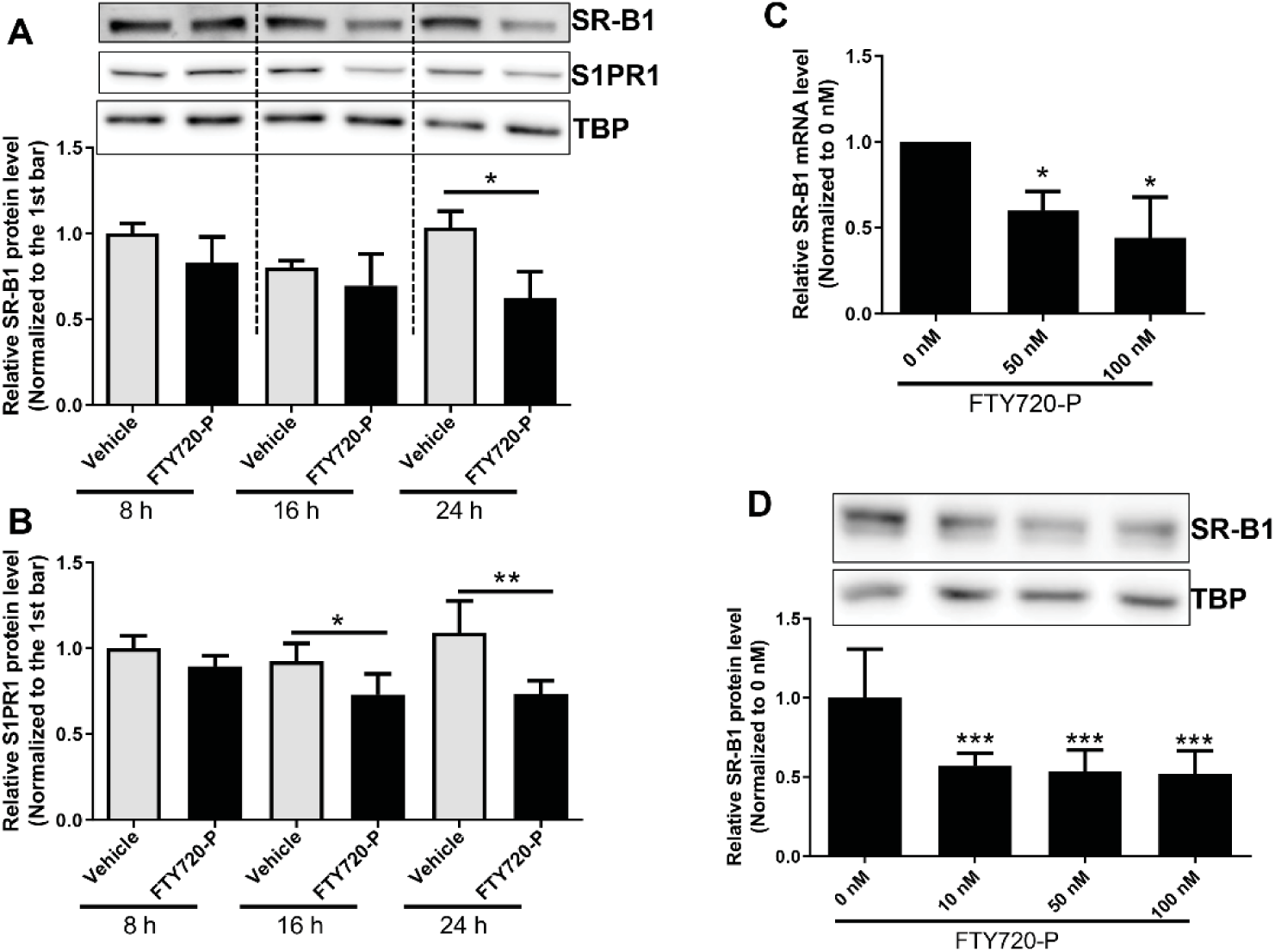
FTY720 phosphate (FTY720-P) decreases both SR-B1 protein and mRNA expression. HAECs were seeded at a density of 0.4 × 10^6^ cells/well in 6-well plates and treated with FTY720-P (**A and B**: 10 nM for 8, 16 or 24 h; **C and D**: 0 - 100 nM for 24 h) or solvent vehicle (0.1% DMSO). The expression of SR-B1 protein and mRNA was determined by western blot analyses and qRT-PCR, respectively. Data are shown as mean ± SD from three independent experiments. *P < 0.05, **P < 0.01, **P < 0.001 (Student’s *t*-test or ANOVA with Bonferroni test).

Taken together the data indicate that both knockdown and pharmacological inhibition of S1PR1 or S1PR3 down-regulate both SR-B1 protein and mRNA levels. Of note, SR-B1 protein expression was suppressed more pronouncedly than *SCARB1* mRNA.

### Decrease of S1PR1 or S1PR3 reduces SR-B1 protein expression on a post-transcriptional level *via* Akt in HAECs

S1PRs were shown to activate various signaling pathways, including Akt (protein kinase B), MEK, p38 MAPK and among others (25). We observed that the knockdown of either S1PR1 or S1PR3 slightly but significantly decreased p-Akt (Ser473) protein levels (Figure 3A). To unravel the signaling pathway by which S1PR1 and S1PR3 regulate SR-B1 expression in HAECs, we used triciribine to inhibit Akt activity and siRNAs against three Akt isoforms as well as siRNAs against MEK or p38 MAPK. Silencing of MEK or p38 MAPK did not influence SR-B1 protein level in HAECs (Supplemental Figure S2). Triciribine (10 and 30 μM) significantly decreased protein level of both phospho-Akt (p-Akt) and SR-B1 by around 65% and 40-45% respectively already after 1h of incubation (Figure 3B). The effect persisted for 8h (Figure 3B). However, triciribine (10 and 30 μM) did not influence *SCARB1* mRNA levels (Figure 3C). Knockdown of Akt1 but not Akt 2 or Akt3 significantly down-regulated SR-B1 protein expression by 44% (20 nM siRNA) (Figure 3D-F). Since Akt is known to activate forkhead box class O family member protein 1 (FoxO1 transcription factors which in turn increases *SCARB1* expression in liver (32), we also tested the effect of FoxO1 knockdown on the expression of *SCARB1*/SR-B1. Neither *SCARB1* mRNA nor SR-B1 protein levels were changed (Supplemental Figure S2C-E). Taken together, the data suggest that Akt1 regulates SR-B1 protein levels post-translationally. We therefore investigated the impact of Akt inhibition by triciribine on SR-B1 protein stability by suppressing *de novo* protein synthesis with cycloheximide (100 μM) and detecting SR-B1 protein levels at different time points (0, 1, 2, 3, 4, and 6 h). The protein half-life (t_1/2_) for SR-B1 in the absence and presence of triciribine were 3.8 and 2.1 h, respectively (Figure 3G). Thus, the decreased SR-B1 protein level induced by triciribine was due to enhanced degradation. Taken together, the data suggest that knockdown of S1PR1 or S1PR3 reduced SR-B1 protein partly by inducing SR-B1 protein degradation through deceasing Akt activity.

**Figure 3.**
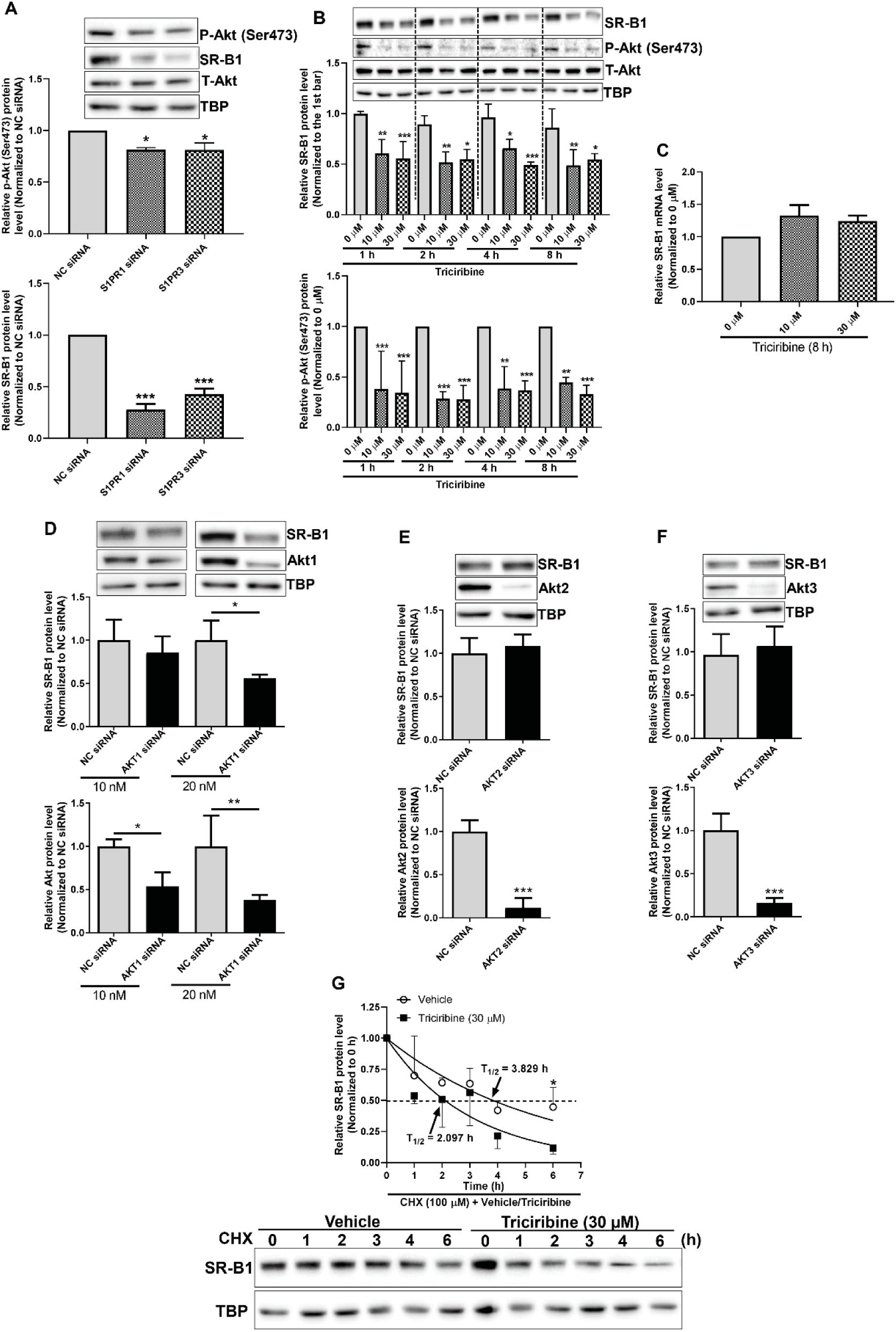
Decrease of S1PR1 or S1PR3 protein expression enhances SR-B1 protein degradation *via* inhibiting Akt activity in HAECs. **A**. S1PR1 or S1PR3 knockdown decreases phospho-Akt (p-Akt (Ser473)) protein level. HAECs were seeded and transfected with siRNA (10 nM) targeted to S1PR1 or S1PR3 as described in **Figure 1A**. The protein expression of SR-B1, p-Akt (Ser473), and total (T-Akt) was determined by western blot analyses. TBP was used as the internal control. **B**. The Akt inhibitor triciribine decreases SR-B1 protein expression along with down-regulation of p-Akt (Ser473) expression. HAECs were seeded at a density of 0.4 × 10^6^ cells/well in 6-well plates for 72 h. The cells were treated with triciribine (0 - 30 μM) for 1, 2, 4 or 8 h. The expression of SR-B1, p-Akt (Ser473), and T-Akt protein was determined by western blot analyses. TBP was used as the internal control. **C**. Triciribine does not influence SR-B1 mRNA level. HAECs were seeded at a density of 0.4 × 10^6^ cells/well in 6-well plates for 72 h. The cells were treated with triciribine (0 - 30 μM) for 8 h. The expression of SR-B1 mRNA was determined by qRT-PCR. GAPDH mRNA was used as the internal control. **D, E and F**. Knockdown of Akt1, but not Akt2 or 3, decreases SR-B1 protein expression. HAECs were seeded and transfected with siRNA targeted to Akt1 (10 and 20 nM), Akt2 (10 nM) or Akt3 (10 nM) as described in **Figure 1A**. The protein expression of SR-B1, Akt1, 2 and 3 was determined by western blot analyses. **G**. Triciribine enhances SR-B1 protein degradation rate. HAECs were seeded as described in **Figure 3A**. Cells were lysed at different time points (0, 1, 2, 3, 4, and 6 h) after incubation with the *de novo* protein synthesis inhibitor cycloheximide (CHX, 100 μM) and triciribine (30 μM) or vehicle (0.1% DMSO). The SR-B1 protein expression was determined by western blot analyses. Data are shown as mean ± SD from three independent experiments. *P < 0.05, **P < 0.01 and **P < 0.001 (Student’s *t*-test or ANOVA with Bonferroni test).

### Silencing of S1PR1 or S1PR3 reduces SR-B1 transcriptional expression *via* cAMP in HAECs

We next started to unravel the mechanism by which knock-down of S1PR1 and S1PR3 suppresses *SCARB1* mRNA expression in HAECs. The established inducers of *SCARB1* gene expression are LXR and RXR and the second messenger cAMP (15). Both the LXR activator TO901017 and the RXR activator bexarotene increased the expression of the positive control gene ABCG1 in HAECs, no matter whether or not S1PR1 or S1PR3 were silenced. However, TO901017 and bexarotene neither altered SR-B1 expression under baseline condition nor rescued the loss of SR-B1 expression after S1PR knockdown (Supplemental Figure S3). However, the exogenous addition of cAMP rescued both SR-B1 protein and *SCARB1* mRNA expression reduced by S1PR1 or S1PR3 knockdown (Figure 4A and 4B). S1PR1 or S1PR3 knockdown reduced cellular levels of cAMP dramatically both in the presence or absence of S1PR1 (SEW2871) or S1PR3 agonists (CYM5541) (Figure 4C). These data indicate that silencing of S1PR1 and S1PR3 regulate *SCARB1* mRNA expression by decreasing cellular cAMP levels.

**Figure 4.**
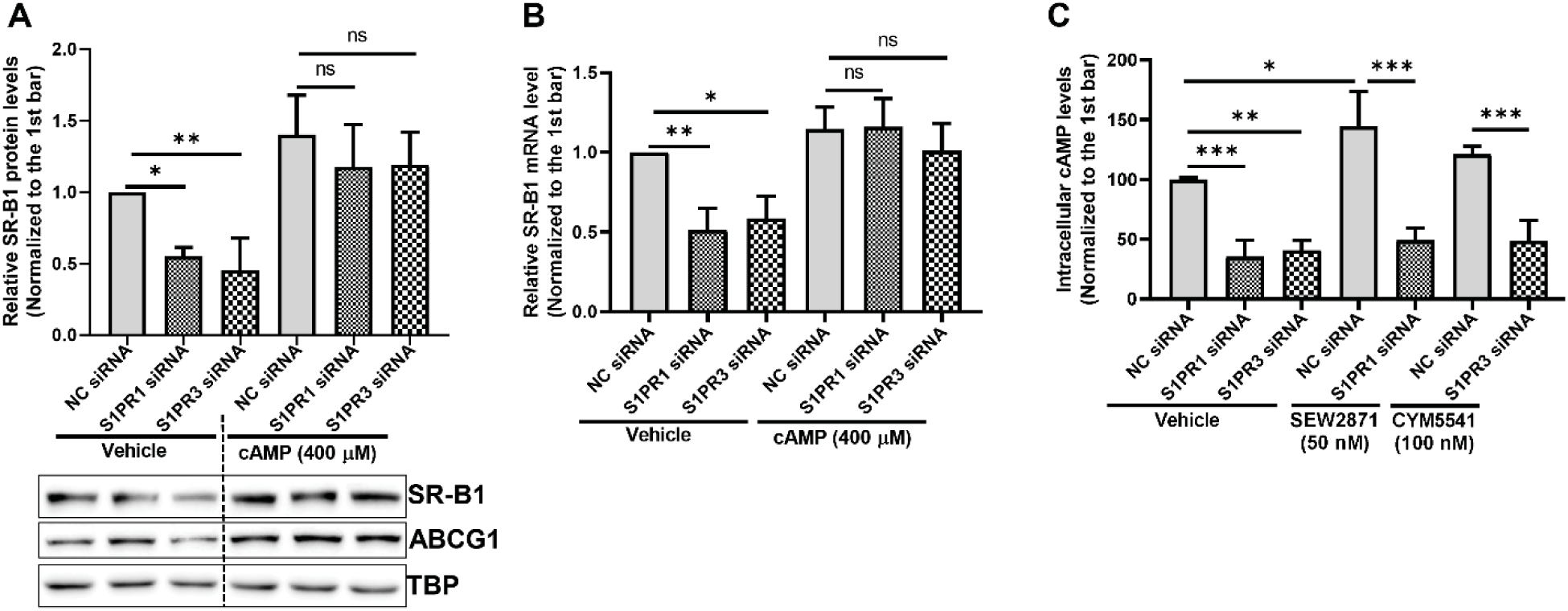
Knockdown of S1PR1 or S1PR3 reduces mRNA expression of *SCARB1 via* cAMP in HAECs. **A and B.** Exogenous addition of cAMP rescues the protein and mRNA levels of SR-B1 reduced by S1PR1 or S1PR3 knockdown. HAECs were seeded and transfected with siRNA (10 nM) targeted to S1PR1 or S1PR3, or with NC siRNA as for 48 h described in **Figure 1A**. The cells were then treated with cAMP (400 μM) or vehicle (0.1% DMSO) for 16 h. The protein and mRNA levels of SR-B1 were determined by western blot analyses and qRT-PCR, respectively. ABCG1 protein expression was used to confirm the positive effect of cAMP. **C**. S1PR1 or S1PR3 knockdown reduces cellular cAMP level in the presence or absence of the S1PR1 (SEW2871) or S1PR3 agonists (CYM5541). HAECs were seeded and transfected with siRNA (10 nM) targeted to S1PR1 or S1PR3, or with NC siRNA as for 72 h described in **Figure 1A**. The cells were then treated with SEW2871 (50 nM), CYM5541 (100 nM) or vehicle (0.1% DMSO) for 1h. The cellular cAMP level was measured by using cAMP-Glo™ Assay kit. Data are shown as mean ± SD from three independent experiments. *P < 0.05, **P < 0.01, ***P < 0.001, and ns not significant (Student’s *t*-test or ANOVA with Bonferroni test).

## DISCUSSION

HDL exerts many vasoprotective functions on the endothelium including stimulation of nitric oxide production and angiogenesis and the maintenance of barrier functions towards leukocytes and macromolecules (33). Several endothelial responses of endothelial cells to HDL have been shown to depend on the presence of S1PRs as well as SR-B1 (18, 20-24). As yet it is not clear whether these receptors act in parallel or in series. A transient triple interaction between HDL, SR-B1, and S1PRs was reported to elicit calcium flux and S1PR1 internalization (34). In this model, SR-B1 facilitates functions of S1P receptors. We here provide evidence for the additional converse relationship, namely that S1PR1 and S1PR3 regulate the abundance of SR-B1 in HAECs: Suppression or pharmacological inhibition of S1PR1 or S1PR3 down-regulates *SCARB1* mRNA expression as well as SR-B1 protein abundance. At least two pathways and mechanisms appear to be involved, one transcriptional involving cAMP and one post-translational involving Akt1 (Figure 5).

**Figure 5.**
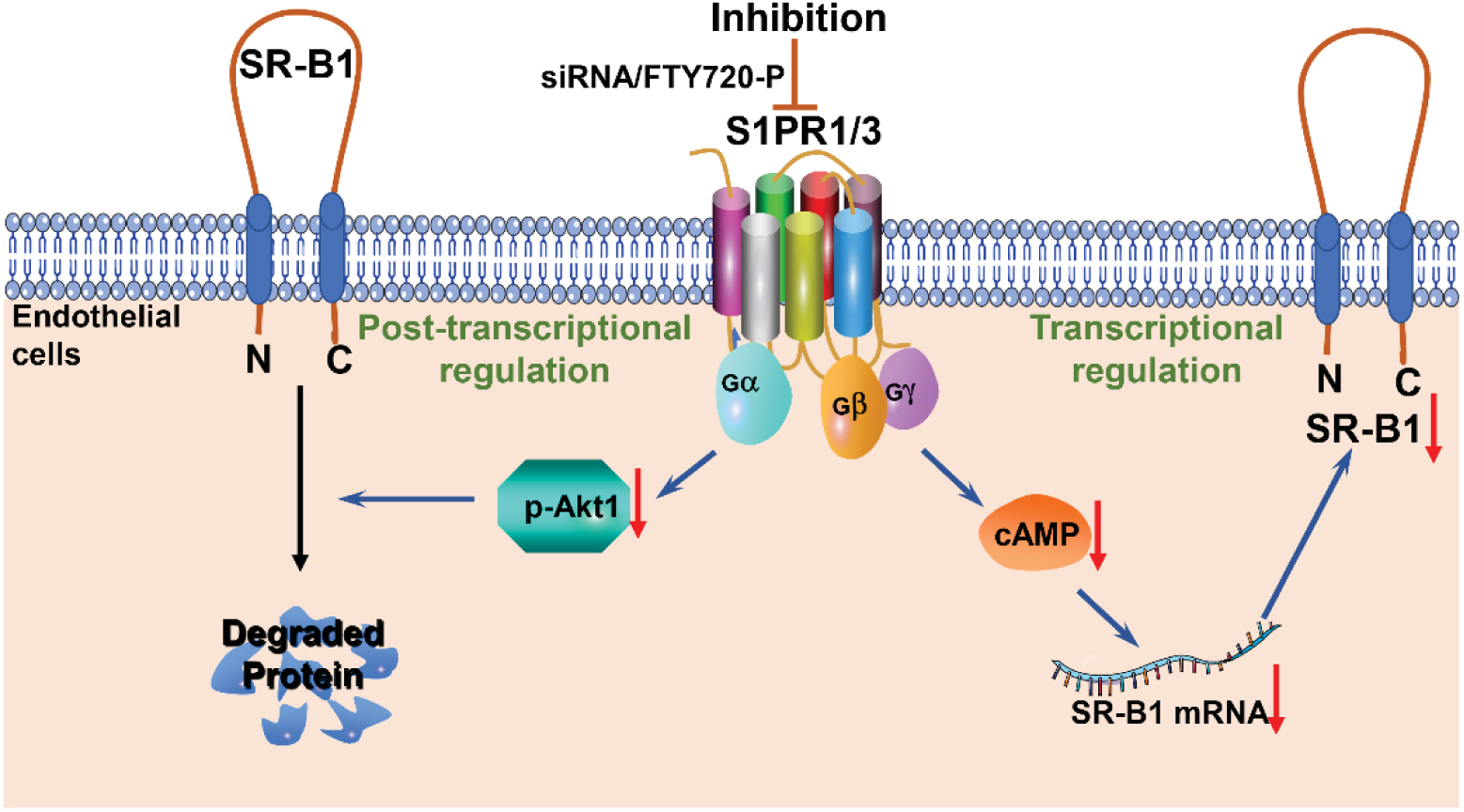
Loss of function of S1PR1 or S1PR3 limits SR-B1 expression in HAECs on the transcriptional and posttranslational levels. First, loss of S1PR1 or S1PR3 activity enhances SR-B1 protein degradation via decreased phosphorylation of Akt1. Second, silencing of S1PR1 or S1PR3 suppresses *SCARB1* mRNA and, as the consequence, SR-B1 protein levels, by a decrease in cAMP.

Endothelial cells express S1PR1, S1PR2, and S1PR3 (21), however to different extent in vessels of various tissues. S1PR2 was found to be the predominant S1P receptor expressed in human coronary artery endothelial cells (35). In our HAEC model, S1PR1 and S1PR3 were prominently expressed on the protein level whereas S1PR2 protein was hardly detectable. The data are in agreement with data of a previous study that found mRNAs of endothelial differentiation gene 1 (EDG1, S1PR1) and EDG2 (S1PR3) but not EDG5 (S1PR2) expressed in HAECs (36). We therefore focused our experiments on S1PR1 and S1PR3. Moreover, it is important to note that previous studies unraveled a specific role of S1PR1 for the endothelial effects of HDL-associated S1P, for example inhibition of VCAM and ICAM1 expression or decreasing permeability for macromolecules (23, 37-39).

In line with the findings of previous studies (35, 40, 41), we observed that silencing of S1PR1 or S1PR3 decreases p-Akt (Ser473) protein expression in HAECs. Akt-phosphorylation is involved in several effects of both S1P and HDL on endothelial cells, namely the stimulation of NO production and angiogenesis as well as inhibition of adhesion molecule expression and apoptosis. All of these effects were also reported to involve SR-B1 (20, 21, 23, 24). We here show that Akt limits the degradation of SR-B1. In the presence of the Akt inhibitor triciribine, the half-life of SR-B1 in HAECs is shortened from about 4h to about 2h. In this regard it is important to refer to another posttranslational effect of Akt on SR-B1: Previous cell surface biotinylation experiments of our lab showed that during 30 minutes of incubation VEGF-A promotes the translocation of SR-B1 from intracellular compartments to the cell surface by an Akt-dependent mechanism (12). Likewise, very short incubation of HAECs with agonists of S1PR1 or S1PR3 promote the translocation of intracellular SR-B1 to the cell surface (*S. Velagapudi, et al*., *submitted for publication*). Probably there are two post-translational pathways, by which Akt regulates the activity of SR-B1 in HAECs, namely the inhibition of degradation and the promotion of cell surface abundance. Akt also regulates the expression of SR-B1 in hepatocytes. In HepG2 cells, like in our HAECs, this happens on the post-translational level (42). However, in mouse liver FoxOs transcription factors, which are activated by Akt, regulate the expression of *SCARB1* (32). Of note, we did not find any involvement of FoxO1 transcription factors in the regulation of *SCARB1* expression in HAECs, neither on the mRNA level nor on the protein level.

Independently of Akt, S1PR1 and S1PR3 also regulate *SCARB1* expression on the transcriptional level. Transcription factors known to induce *SCARB1* expression in different cell types include FoxOs (32) and cAMP response-element binding protein (CREB) (43). As mentioned before, we excluded any involvement of FoxO transcription factors in the regulation of *SCARB1* in HAECs. Likewise and in contrast to previous findings in human umbilical vein endothelial cells (44), in our HAEC model, TO901317 and bexarotene did not increase the expression *SCARB1* mRNA. Our data however suggest that the transcriptional regulation involves cAMP because RNA interference with S1PR1 or S1PR3 decreased cellular cAMP levels and because the suppression of *SCARB1* by interference with S1PR1 or S1PR3 was prevented by the addition of exogenous cAMP. In agreement with our findings, cAMP was reported to increase *SCARB1* mRNA expression in transfected 293T cells (43), rat theca-interstitial cells (43) and adrenocortical cells (45). Moreover, in human coronary artery smooth muscle cells, S1P was shown to elicit cAMP accumulation, in a S1PR2 dependent manner (46). It remains to be unraveled how S1PR1/3 knockdown regulates cAMP level in HAECs, by stimulating an adenylate cyclase or inhibiting a phosphodiesterase.

Both SR-B1 and S1PRs contribute to protective effects of HDL on the endothelium (47). However, SR-BI was also found to mediate transendothelial transport of both HDL and pro-atherogenic LDL (9-13). Also of note, the transendothelial transport of HDL and LDL is regulated by S1P3 into opposite directions: Activation and overexpression of S1PR3 in endothelial cells, promote the tranendothelial transport of HDL but inhibit the transendothelial transport of LDL in vitro and in vivo, respectively (*Velagapudi, Wang, Poti et al. submitted for publication*). Both endothelium-specific knock-out of *Scarb1* and overexpression of SCARB1 was reported to reduce atherosclerosis in mice (9,48). Likewise controversial data has been reported on the impact of modulated S1P metabolism or S1P receptor activity on atherosclerosis (49-54). The majority of data rather point to protective effects but none of them specifically investigated the endothelium-specific effects of S1P on atherosclerosis.

In conclusion, we here provide evidence for an as yet unappreciated interaction, namely the regulation of SR-B1 abundance by S1PRs on both transcriptional and post-translational levels. Together with previous observations indicating that SR-B1 facilitates S1PR activation, our findings help to solve the controversy on the role of S1P and its G-protein coupled receptors versus SR-B1 in mediating protective functions of HDL on the vasculature as well as transendothelial transport of HDL and LDL.

## Abbreviations

Akt: protein kinase B;
cAMP: cyclic AMP;
HAEC: human aortic endothelial cells;
HDL: high-density lipoprotein;
LDL: low-density lipoprotein;
LXR: liver X receptor;
MEK: extracellular signal-regulated kinase kinase;
p38 MAPK: p38 mitogen-activated protein kinase;
RLU: relative luminescence;
RXR: retinoid X receptor;
S1P: Sphingosine-1-phosphate;
S1PR: Sphingosine-1-phosphate receptor;
siRNA: small interfering RNA;
SR-B1: scavenger receptor class B type 1;

## Data availability statement

All data are contained within the manuscript and supplemental data file.

## Acknowledgments/grant support

This work was supported by grants from the Peter und Traudl Engelhorn Foundation for the promotion of Life Sciences to D.W. and the Swiss National Science Foundation (31003A-160216 and 310030_166391/1) to A.v.E.

## Figures and figure legends

**Supplemental Figure S1.**
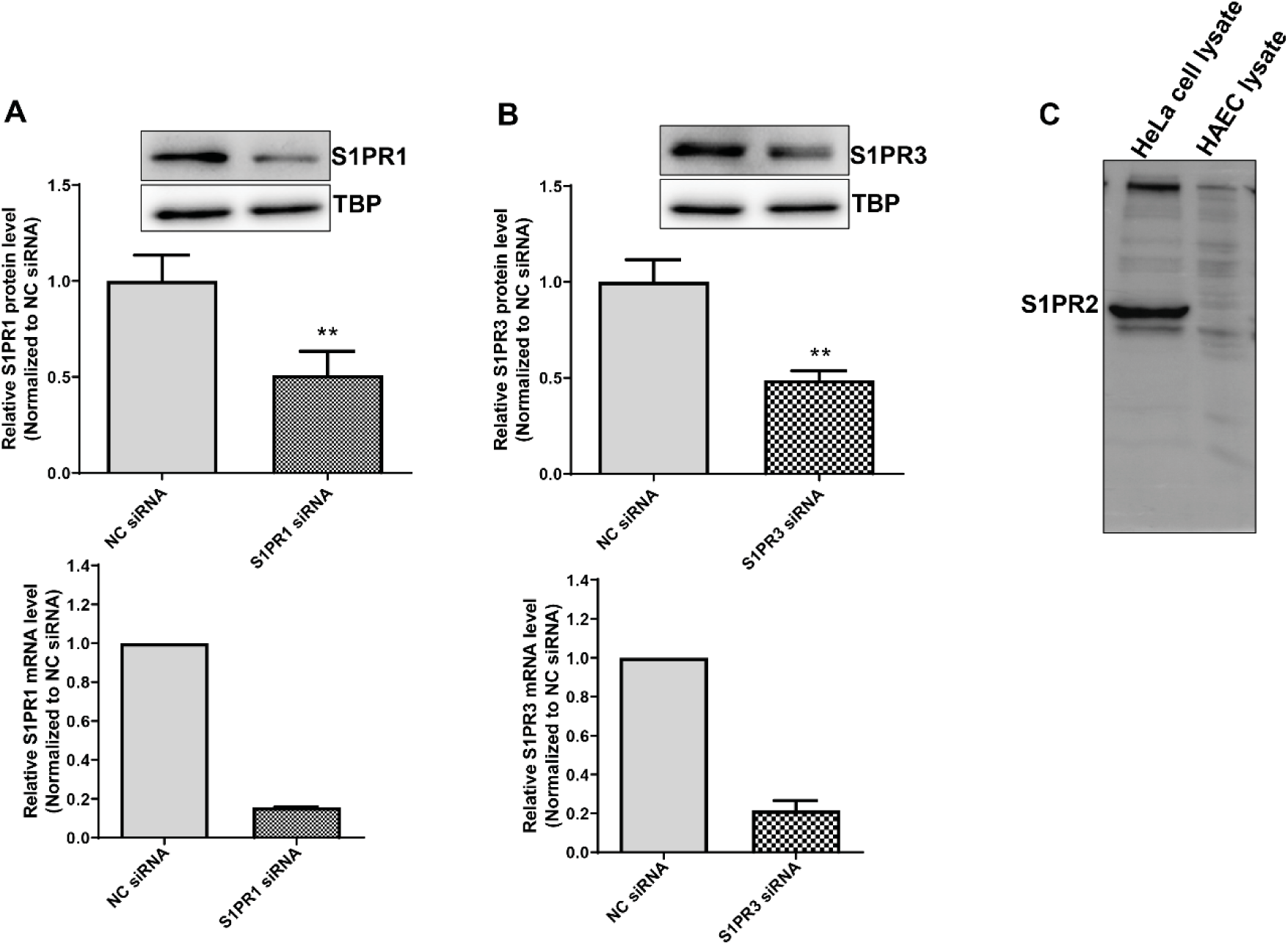
A and B. Knockdown efficiency of S1PR1 and S1PR3 in HAECs. HAECs were seeded at a density of 0.4 × 10^6^ cells/well in 6-well plates. The cells were reverse transfected with siRNA (10 nM) targeted to SR-B1, S1PR1 or S1PR3, or with non-silencing control siRNA (NC siRNA) for 72 h. The expression of SR-B1 protein and mRNA was determined by western blot analyses and RT-qPCR, respectively. TATA-binding protein (TBP) and GAPDH mRNA were used as the internal control for western blot analyses and RT-qPCR, respectively. Data are shown as mean ± SD from three independent experiments. **P < 0.01 (Student’s *t*-test). **C**. Low expression of S1PR2 protein in HAECs. Hela cell lysate is as a positive control, which over-expressed S1PR2 protein.

**Supplemental Figure 2S.**
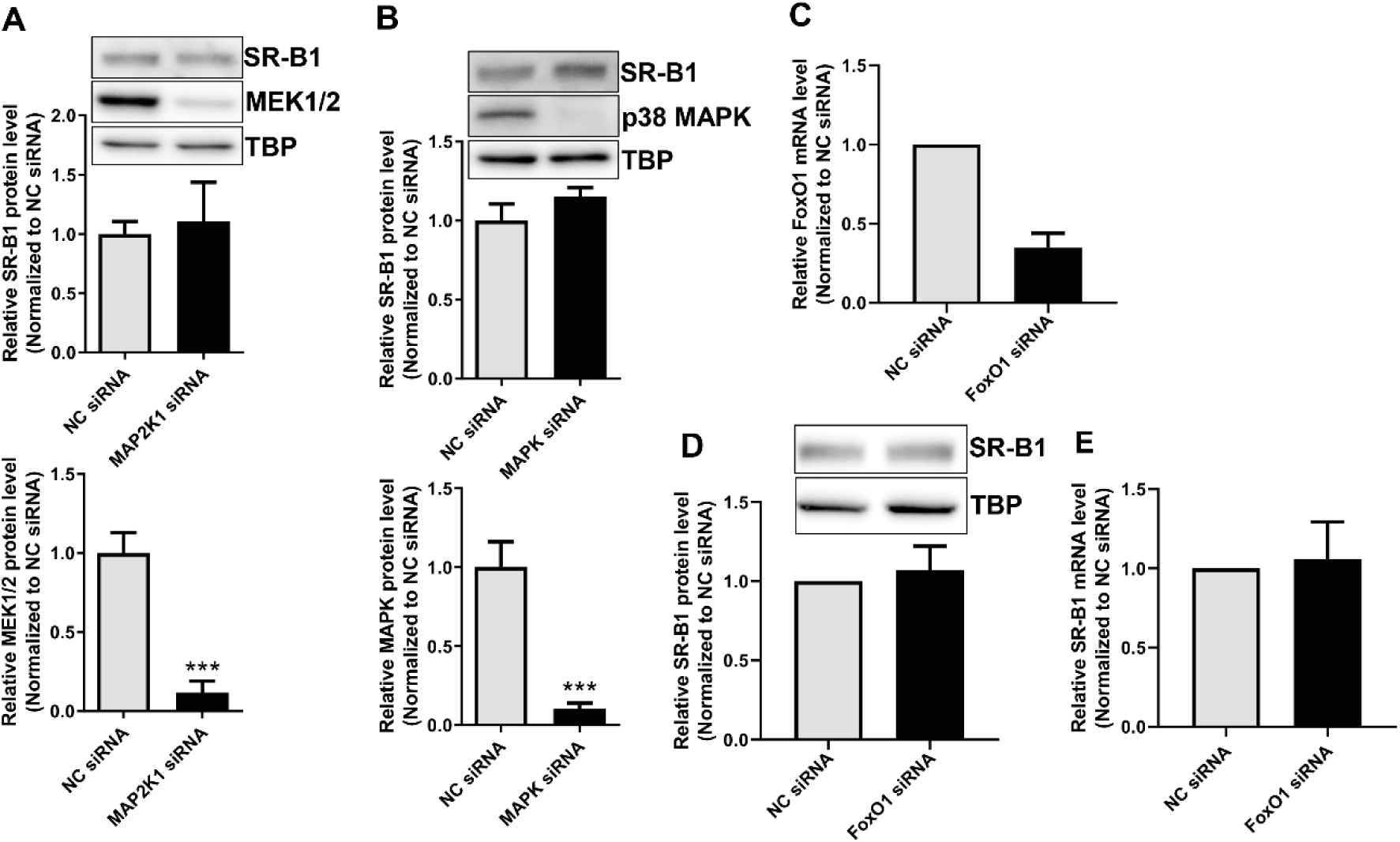
**A and B. Knockdown of MEK1/2 (A) or p38 MAPK (B) does not influence SR-B1 protein expression in HAECs.** HAECs were seeded and transfected with siRNA (10 nM) targeted to MEK1/2, or p38 MAPK as described in **Figure 1S**. The protein expression of SR-B1, MEK1/2 and p38 MAPK was determined by western blot analyses. TBP was used as the internal control. The data of samples were normalized to NC siRNA. **C-E. Knockdown of FoxO1 does not influence SR-B1 mRNA and protein expression in HAECs**. HAECs were seeded and transfected with siRNA (10 nM) targeted to FoxO1 as described in **Figure 1S**. The protein expression of FoxO1 was determined by western blot analyses. TBP was used as the internal control. The data of samples were normalized to NC siRNA. Data are shown as mean ± SD from three independent experiments. ***P < 0.001 (Student’s *t*-test).

**Supplemental Figure S3.**
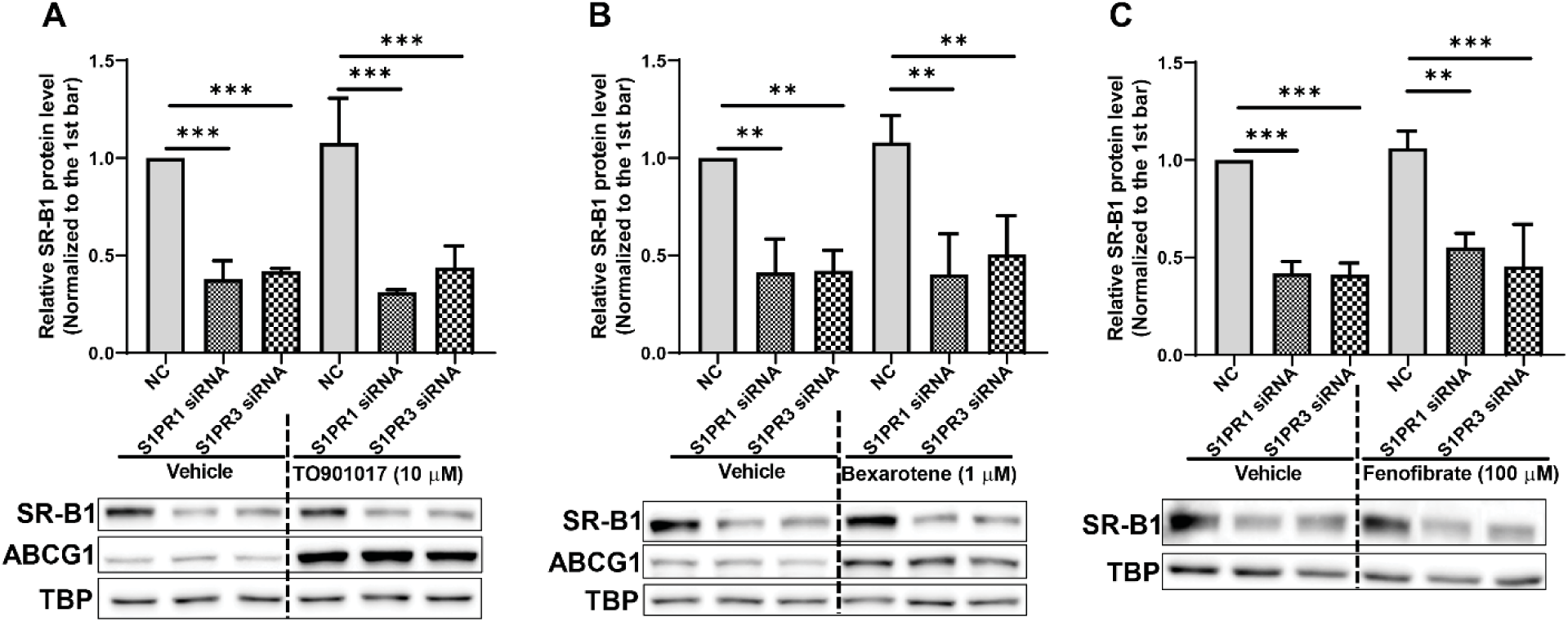
Activation of LXR or RXR does not restore SR-B1 protein level reduced by S1PR1 or S1PR3 knockdown. HAECs were seeded and transfected with siRNA (10 nM) targeted to S1PR1 or S1PR3, or with NC siRNA as for 48 h described in **Figure S1**. The cells were then treated with the LXR agonist TO901017, RXR agonist bexarotene, or vehicle (0.1% DMSO) for 16 h. The SR-B1 and ABCG1 protein expression were determined by western blot analyses. ABCG1 protein expression was used to confirm the positive effect of TO901017 and bexarotene. Data are shown as mean ± SD from three independent experiments. **P < 0.01, and ***P < 0.001 (Student’s *t*-test or ANOVA with Bonferroni test).

